# Plant-herbivore interactions: experimental demonstration of genetic variability in plant-plant signaling

**DOI:** 10.1101/2022.06.28.497952

**Authors:** Aurélien Estarague, Cyrille Violle, Denis Vile, Anaïs Hany, Thibault Martino, Pierre Moulin, François Vasseur

## Abstract

Plant-herbivore interactions mediated by plant-plant signaling were documented in different species. Here, we tested if herbivore foraging activity on plants was influenced by plant’s prior contact with a damaged plant and if the effect of such plant-plant signaling was variable across genotypes.
We filmed snails during one hour on two plants differing only in a prior contact with a damaged plant or not. We replicated eight times the experiment on 113 natural genotypes of *Arabidopsis thaliana*. We recorded snails’ first choice, and measured its first duration on a plant, the proportion of time spent on both plants and leaf consumption.
On average, snails spent more time on plants that experienced a prior contact with a damaged plant, and consumed them more. However, plant-plant signaling effect on snail behavior was variable across genotypes. Genome-wide association studies revealed that a small number of genetic polymorphisms related to stress coping ability and jasmonate pathway explained this variation.
Plant-plant signaling modified the foraging activity of herbivores in *A. thaliana*. Depending on the plant genotype, plant-plant signaling made undamaged plants more repulsive or attractive to snails. This finding questions the theoretical basement of the evolution of plant-herbivore interactions mediated by plant-plant signaling.

**Highlight:** Plant-plant signaling differently affects snail foraging activity depending on genetic variations in *A*.*thaliana*. These findings question the theoretical basement of the evolution of plant-herbivore interactions mediated by plant-plant signaling.

## Introduction

Plants adjust their defenses in response to cues from neighbors that are being attacked by herbivores (Agrawal, 2000; Karban et al., 2000, 2003; Heil and Adame-Álvarez, 2010; Heil and Karban, 2010; Moreira et al., 2018; Bouwmeester et al., 2019; Moreira and Abdala-Roberts, 2019). Interactions between damaged plants and their neighbors are generally associated with increased plant defense level. This phenomenon appears widespread across biomes, growth forms and phylogeny (Karban et al., 2014; Karban, 2021). Such plant-herbivore interactions mediated by plant-plant signaling is often considered adaptive, as it reduces plant biomass loss and then supposedly increases individual fitness (Heil and Karban, 2010). This suggests that plant-plant signaling could be variable within species and differentiate between populations as the result of herbivory pressure and natural selection.

Plant-plant signaling can have prompt effects on herbivore foraging activity. A contact with a damaged plant can prime defenses, expressed then faster in case of an herbivore attack (Engelberth et al., 2004; Heil and Bueno, 2007; Zhang et al., 2020). Plant-plant signaling can also directly trigger neighbor defenses, that take place at different steps of the foraging activity of surrounding herbivores. Plants that receive herbivore-induced signals emitted by a neighbor can express volatile molecules, that constitute a distant and repellent signal for herbivores (Engelberth et al., 2004; Shannon et al., 2016; Morrell and Kessler, 2017). Receivers of herbivore-induced signals can also produce secondary metabolites, or toxins within leaves (Himanen et al., 2010; Sugimoto et al., 2014) that limit the palatability of receiver plants when herbivores feed on them. Leaf consumption on receiver plants of herbivore-induced signals was mainly measured over long period of time (days, weeks or years) (Rhoades, 1983; Fowler and Lawton, 1985; Agrawal, 2000; Karban et al., 2000, 2003, 2016; Pearse et al., 2013; Morrell and Kessler, 2017; Benevenuto et al., 2020). Recent studies also showed an effect of plant-plant signaling on herbivore fitness component, as their fecundity or their growth (Morrell and Kessler, 2017; Moreira et al., 2018). Understanding real-time foraging activity of herbivores on plants may be the key to understand the settings of plant defenses following a prior contact with a damaged plant (Morrell and Kessler, 2017).

Genetic variability in plant-plant signaling impacting herbivore activity is a prerequisite condition for natural selection (Falconer et al., 1996). Many field and lab experiments suggest high genetic variability on plant-herbivore interactions mediated by plant-plant signaling. Both nature and concentration of volatiles emitted after herbivore damages are highly variable within and among species (Chen et al., 2003; Degen et al., 2004; Dudareva et al., 2006; Snoeren et al., 2010; Kalske et al., 2019; Müller et al., 2020). Plant-plant signaling can evolve in a few generations only, depending on the herbivory pressure existing within populations (Kalske et al., 2019). For instance, environmental clines, such as altitudinal gradients, has a strong effect on plant-herbivores interactions mediated by plant-plant signaling (Benevenuto et al., 2020). In their study, Benevenuto and colleagues (2020) showed that simulated attacks of herbivore on bilberry plants reduced leaf consumption on conspecific neighbors at low and mid altitude, but had no effect at high altitude, indicating potential costs of induced defenses in stressful environments. Here, we used a highly cosmopolitan plant, *Arabidopsis thaliana*, to investigate the potential intraspecific variability of the effect of plant-plant signaling on plant-herbivore between genotypes originating from various environmental conditions.

*Arabidopsis thaliana* L. is a small annual plant inhabiting contrasted environments and climates all over the world (Hoffmann, 2002; 1001 Genomes Consortium, 2016; Lee et al., 2017). An international effort of sampling made available fully-sequenced genomes for hundreds natural genotypes (hereafter ‘accessions’) (1001 Genomes Consortium, 2016). This open-access database is a great opportunity to study the genetic bases of phenotypic variation. For instance, genomic variations between accessions of *A. thaliana* showed evidence of local adaptations to climate (Lasky et al., 2014; Dittberner et al., 2018; Vasseur et al., 2018; Exposito-Alonso, 2020; Clauw et al., 2022). Moreover, natural accessions of *A. thaliana* expressed contrasted values of constitutive defenses against herbivores, notably in leaf glucosinolate concentrations (Brachi et al., 2015; Kerwin et al., 2015; Gloss et al., 2017). Snoeren et al (2010) showed that nine accessions of *A. thaliana* emitted different blends of volatile molecules consecutively to herbivore damages (see also Savchenko et al., 2013). Other studies showed that Col-0 accession increased gene expression associated with resistance functions after being exposed to particular volatile molecule such as aldehyde or monoterpenes (Kishimoto et al., 2005; Savchenko et al., 2013). Surprisingly though, the effect of a prior contact with herbivore-induced signal on the foraging activity of an herbivore have been poorly investigated in this model species. The natural variation of the response of such herbivore-induced volatiles within *A. thaliana* is also an opportunity to understand its genetic bases through genome-wide association studies (GWAS). Such approaches are powerful tools to link complex phenotypes to genetic variation (Francisco et al., 2016).

Here we asked: (i) Is the foraging activity of a generalist herbivore impacted by the prior contact of a plant with another damaged plant? (ii) Does plant-herbivore interaction mediated by a prior contact vary across natural genotypes of *Arabidopsis thaliana*? (iii) Is this potential variation explained by allelic diversity between plant genotypes? To address these questions, we conducted a glasshouse experiment with 113 worldwide accessions of *A. thaliana*. We built an innovative experimental design (**Figure 1**) that filmed the foraging activity of snails (*Helix aspera*) across pairs of plants differing only by a prior contact or not with an artificially damaged individual.

**Figure 1:**
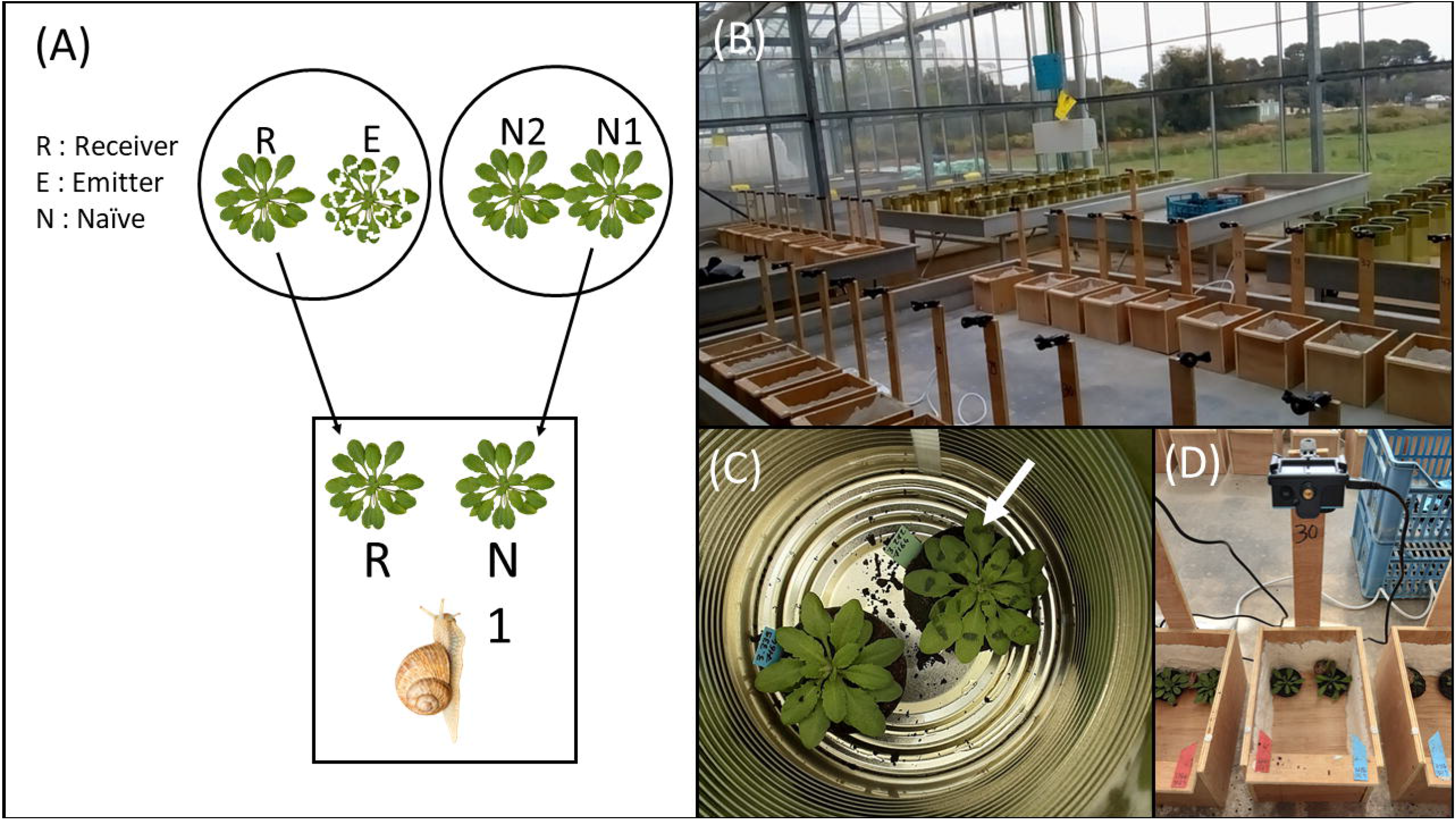
Experimental test of the effect of a prior contact with a damaged plant on the foraging activity of snails. (A) scheme of the two stages of the experiment. (B) Picture of the greenhouse and of the experimental devices. (C) Details of a circular metallic enclosure. The contact between an emitter plant E (noted with an arrow, clipped) and a receiver plant R lasted 30 minutes in parallel of two intact plants (N1 and N2). (D) Woody arena where snail movements on R and N1 were filmed during 1h.

## Material and Methods

### Plant material

We randomly selected, in our in-house collection (CEFE, Montpellier, France), 113 natural genotypes (accessions) of *Arabidopsis thaliana* from Eurasia. All the accessions were initially obtained from the 1001 Genome germplasm and were entirely sequenced (6,973,565 single-nucleotid polymorphisms, SNPs (1001 Genomes Consortium, 2016)). We used *Helix aspera*, a generalist herbivore, whose geographical range overlap the quasi-totality of the European distribution of *Arabidopsis thaliana*. Snails used in this study were raised in an organic farm (L’escargot du Riberal, L. Sanchez, Tautavel, France). Snails were fed with industrial and specialized flour at the farm and were starved during one week in a cold room before the experiment.

### Growth conditions

Seeds were sown in October 2020 in a greenhouse at Montpellier, France (CEFE CNRS). We used peat soil (Neuhaus N2) in 5-cm-diameter pots. We realized eight consecutive sowing consisting in four seeds of 113 accessions (*n* = 3616 pots in total), randomly arranged in eight blocks, one block corresponding to one sowing day. All blocks were rotated on a daily basis. All pots were sub-irrigated with water treated by reverse osmosis to field-capacity every two to four days. The temperature was set at 20 °C during the day and 16 °C during the night for the full duration of the experiment. Sciarid flies were observed five weeks after germination. We spread water-diluted larvicide on the surface of all the pots (Vectobac WG, Edialux, France). Due to germination issues and on sciarid flies, 600 pots were removed. The 3016 remaining pots were undamaged by sciarid flies and represented between 20 to 32 individuals per accession.

### Experimental design

Seven weeks after sowing, two pairs of individual plants of the same accession and from the same block were separately placed in circular metallic enclosures (diameter = 15 cm, height = 25 cm, **Figure 1AB**). In the first pair, all leaves of one individual (‘emitter plant’, E) were clipped with clamps in order to simulate herbivory (**Figure 1C**). The neighbor individual (‘receiver plant’, R) was kept intact. Both control individuals of the second pair were left undamaged (‘naive plants’ N1 and N2). The two pairs of neighbor plants remained 30 minutes in their respective enclosure after the treatment of emitter plant E. Then, the intact individuals from the two pairs (R and N1) were placed together in a woody arena (**Figure 1D**). A snail (*Helix aspera*) was released at equidistance of each plant and was filmed in the arena during one hour (1 photo per second, 2.7 kppi, XPRO2, TecTecTec, France). An example of a video is available in the **Supplementary Video S1**. We took pictures of the plants before and after the passage of the snail (16 Gpi, XPRO2, TecTecTec, France). We performed simultaneously 63 tests on a half day, one test corresponding to one replicate per genotype. Positions of pairs of plants among the 126 metallic enclosures and among the 63 arenas were randomly assessed. All videos and images are available at https://doi.org/10.57745/QN7MMM.

### Measurements

A binomial variable ‘First choice’ was scored 1 to every arena where R plant was chosen first by the snail, and 0 elsewhere, *i*.*e*. to every arena where N1 plant was chosen first (**Table 1**). First duration was measured as the time spent by the snail on the first plant (**Table 1**). We measured the total duration that the snail spent on each individual during one hour. The proportion of time was then calculated as the duration on one plant divided by the sum of the durations on the two plants in the arena (**Table 1**). We estimated a ‘leaf consumption index’ from the pictures taken after the passage of the snail (**Table 1**). We analyzed individual pictures (and not pictures of pairs) in order to avoid biased results related to the leaf consumption index of the second individual of the pair. A value of zero was given if the snail did not consume any part of the plant and a maximum value of five was given if all the leaves of the plant were eaten by the snail (**Supplementary Fig. S1**). We confronted the leaf consumption index estimated by two independent observers on a training data set of 227 pictures to validate the method (ρ = 0.86, *P* < 0.0001). Initial rosette area of each plant was estimated from the picture taken before the passage of the snails (ImageJ; version 1.53c, Schneider et al., 2012). We then calculated the ‘rosette area difference’, as the difference between initial rosette areas of the R plant and the N1 plant in each arena.

**Table 1:**
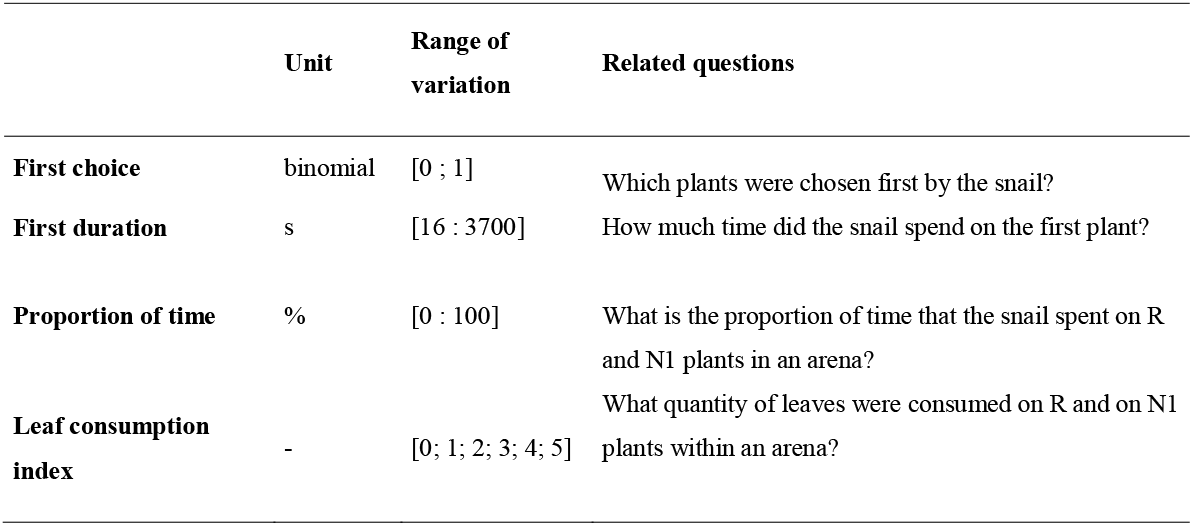
List of response variables and their observed range of variation.

### Statistical analyses

All analyses were performed with R (version 4.0.5). We tested in the same model if a prior contact to a clipped plant and rosette area difference had an effect on the four response variables. We used quasibinomial models to test these effects on the first choice, the proportion of time spent on R, and the proportion of leaf consumption index (package ‘lme4’, link function ‘probit’). We used quasibinomial models instead of binomial model to correct for overdispersion. Quasibinomial models artificially increase standard errors, and are more conservative than binomial models (Dunn and Smyth, 2018). Effect on first duration was tested using a linear model, after log_10_-transformation to fit normality assumptions. Intercepts and slopes estimated by models were used to quantify the effects of prior contact and total leaf area differences between plants, respectively. For example, for the variable ‘First choice’, a significant negative intercept indicates a weaker probability for R plants to be chosen first than N1 plants. A positive slope indicates a higher probability to choose R first than N1 if R plant was bigger than N1 plants.

The effect of accession identity was tested with linear model for the first duration and with quasibinomial models for the three other response variables. Rosette area difference was systematically added as a covariate in these models. The effect of accession identity was reported as its global effect on the variance (anova of type II, package ‘car’). From these models, we calculated marginal means of response variable of all accessions using the package “emmeans” (Lenth et al., 2018). Correlations between mean responses of accessions were assessed with the Kendal’s statistic.

### Quantitative genetic analysis

We tested for allelic effects on the four response variables through genome-wide association studies using the online platform EasyGWAS (https://easygwas.ethz.ch/, Grimm et al., 2017). Marginal means of accessions, as calculated above, were implemented. We used the EMMAX algorithm, filtering for minimum alleles frequency at 5% and including the two first axes of a PCA performed on SNPs to correct for population structure. Significance threshold was assessed by the Bonferroni method with an error alpha of 0.05 and 0.1. Known functions of genes were extracted from TAIR10 database (https://www.arabidopsis.org, Lamesch et al., 2012).

## Results

### Prior exposition to a damaged neighbor and rosette area difference effects on snail foraging activity

Across all accessions, prior contact and rosette area difference did not have a significant effect on the first choice of the snails (*P* = 0.50 and 0.79 respectively, **Figure 2**) nor on the first duration spent on the first encountered individual (*P* = 0.61 and 0.41 respectively, **Figure 2**). The mean first duration on a plant was positively correlated with the mean rosette area (rho = 0.3, *P* = 0.001).

**Figure 2.**
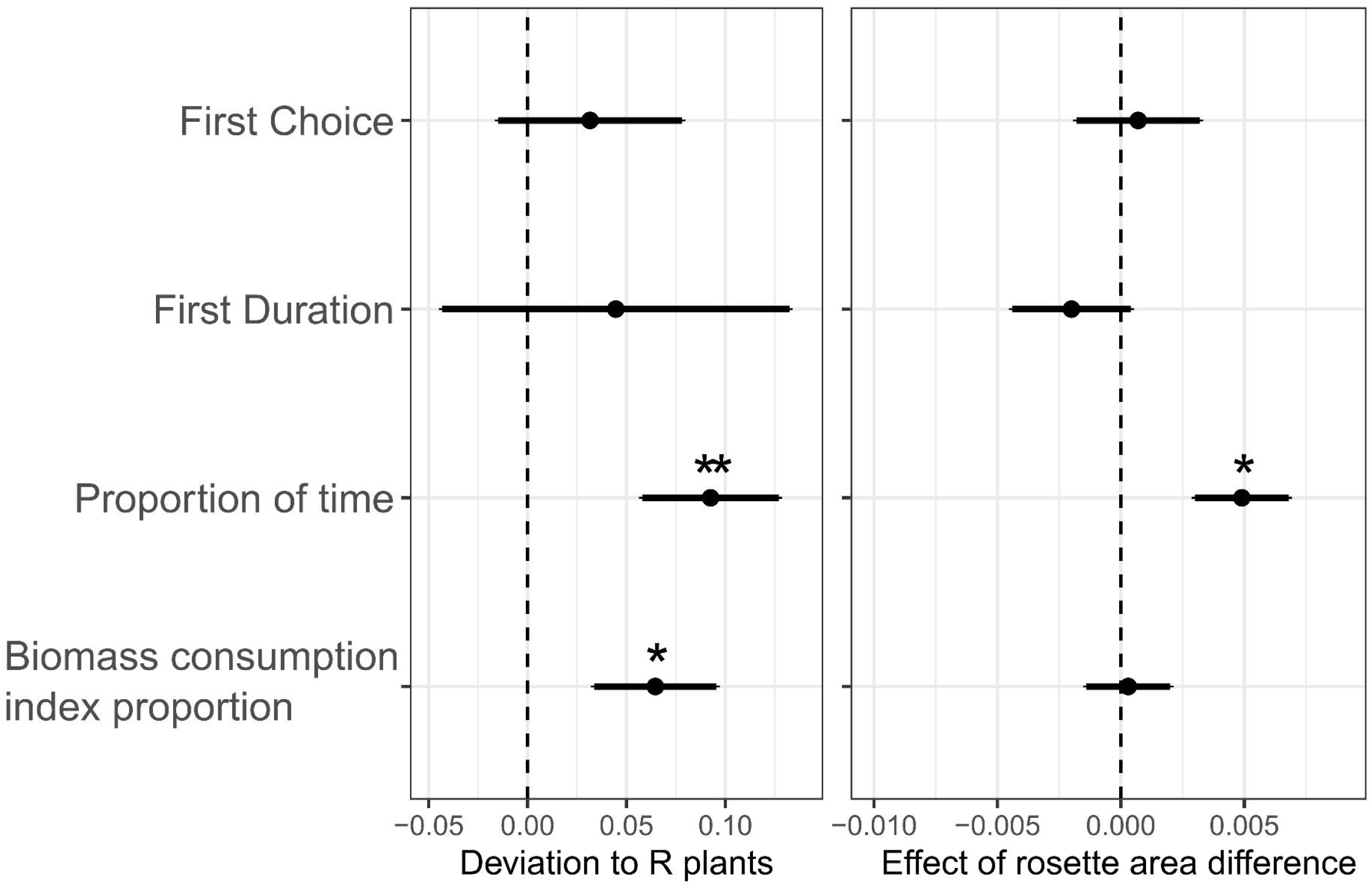
Prior contact and rosette area difference effects on snail foraging activity. Estimates and standard errors of the coefficients of models are represented here on a probit scale. *: *P* < 0.05, **: *P* < 0.01; ***: *P* < 0.001

Prior contact with a damaged neighbor (R plants) had a significant effect on the proportion of time spent on plants (*P* < 0.001). Over all accessions, snails spent proportionally more time on R plants (54%), i.e. plants previously exposed to artificially damaged plants, than on control N1 plants (46%), i.e. plants that were previously adjacent to undamaged plants (**Figure 2**). In addition, difference in rosette area between the two plants in the arena had also a significant and positive effect on the proportion of time spent on R plants (*P* < 0.001, **Figure 2**).

A prior exposition to a damaged neighbor plant had a significant effect on leaf consumption index. Leaf consumption index was higher for R plants than for N1 plants (*P* < 0.05, **Figure 2**). Leaf consumption index was highly correlated with the proportion of time spent on plants (ρ = 0.46, P < 0.001), but not significantly explained by rosette area difference (*P* = 0.88, **Figure 2**).

### Genotypic and allelic effects of attractivity of plants exposed to a damaged neighbor

The probability to choose R plants first varied a lot between plant genotypes, ranging from 12.5% (1/8) to 100% (8/8) across accessions. Two SNPs were associated with first choice of snails on R plants (**Figure 3**). No gene was found at the first SNP on the second chromosome, but this SNP was upstream to AT2G19800, a gene that encodes a myo-inositol oxygenase. On the fourth chromosome, a significant SNP was located in AT4G27080, a gene that encodes a protein disulfide isomerase-like (PDIL) protein (**Supplementary Tab. S1**).

**Figure 3.**
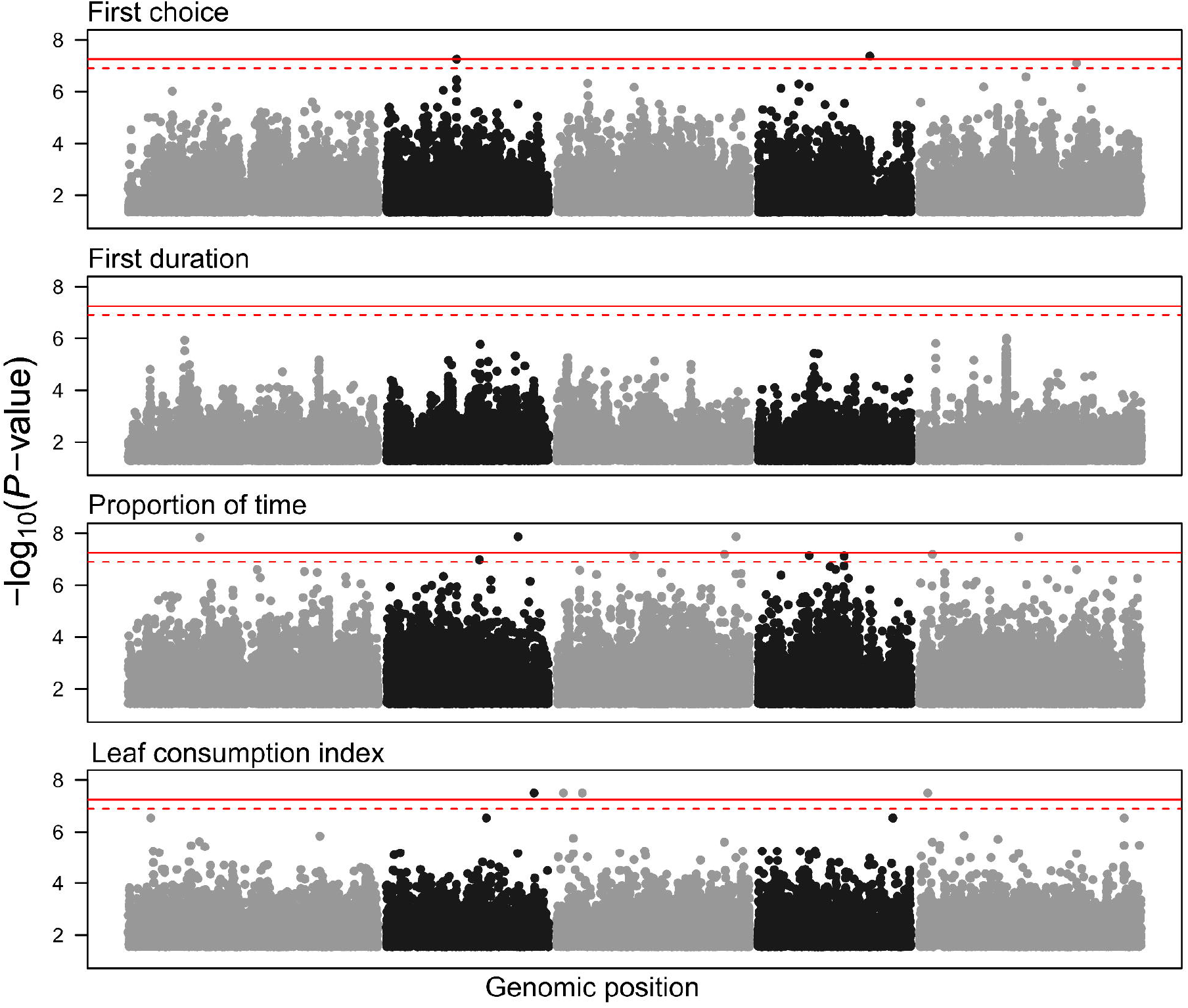
Genome-wide association of the four response variables. GWAS were performed with EMMAX across 113 natural accessions on 1,922,479 SNPs. Each dot is a SNP located on chromosomes 1, 3 and 5 (grey) and chromosomes 2 and 4 (black). Bonferroni threshold at 0.05 and 0.1 was represented with red solid and dashed lines, respectively.

First duration exhibited a bimodal distribution, reflecting that, depending on the accession, the time spent by a snail on the first plant chosen was brief (16.2 to 500 seconds) or long (more than 2000 seconds), with fewer intermediate durations (**Supplementary Fig. S2**). Identity of the accession explained a significant part of the variance of the first duration spent on a plant (*P* = 0.02). However, no SNP position along the genome was significantly associated with first duration spent on a plant (**Figure 3, Supplementary Tab. S2**).

A large variation in the proportion of time spent on R plants across accessions (16% to 100%) was found. Four SNPs along the genomes were significantly associated with the proportion of time spent on R plants (**Figure 3**). Among them, the gene AT1G24560.1 encodes for plant-unique RAB5 effector 2 (PUF2), a key protein in cell-to-cell communication in plants (**Supplementary Tab. S3**).

The leaf consumption index on R plants relative to N1 plants was significantly explained by accession identity (*P* < 0.0001). In other words, depending on the accession identity, a prior exposition to a damaged plant can increase or decrease attractivity of undamaged plants. Along the genome, four SNPs were significantly associated to the proportion of leaf consumption index on R plants (**Figure 3**). Among them, AT3G10050.1 encodes for the first enzyme in the biosynthetic pathway of isoleucine. Another gene was found on chromosome 3 (AT3G03380.1) that encodes for a periplasmic protease (**Supplementary Tab. S4**). The effects and the frequencies of minor alleles for every significant SNP are detailed in **Supplementary Tab. S5**.

## Discussion

For the last decades, consistent progresses have been made in the characterization of plant-plant signaling and its effect on herbivore foraging activity. Within-species variation have been however overlooked, despite its central place in local adaptation. Here we showed that a prior contact with a damaged plant had, on average, a negative effect on *Arabidopsis thaliana’s* defense against a generalist herbivore, as snails spent significantly more time and ate more on receiver plants. However, we also found that this response is very variable between genotypes, and that allelic diversity underlies this variation.

A contact with a damaged plant led to higher attractivity of *A. thaliana* for snails, in terms of time spent on plants and on leaf consumption. Our results confirmed that plant-herbivore interactions can be partly regulated by plant-plant signaling (Dicke and Baldwin, 2010; Karban et al., 2014). However, our results contradicted the general idea that plant-plant signaling has been selected to promote deterrence to herbivore attack (see Pearse and Karban, 2013; Karban et al., 2014; Karban, 2021 for reviews). Snails may be attracted to plants previously exposed to damaged neighbors for many reasons as developed in the followings.

First, odor and taste of the emitter damaged plant can have been deposited on the neighboring plants (Matsui, 2016). Snails possess the ability to sense odor and to discriminate them in a saturated context (Shannon et al., 2016). In the arena, the plant previously exposed to a damaged neighbor, i.e. receiver plant, can thus exhibit a different olfactory signal compared to the plant exposed to an undamaged neighbor. To test the hypothesis that damaged plants can attract snails, damaged and undamaged plants of eleven natural accessions were proposed to snails. Snails significantly preferred damaged plants than undamaged plants (**Supplementary information 1**). Attractiveness of damaged plants was shown in numerous species of insect herbivores (Harari et al., 1994; Loughrin et al., 1995; Bolter et al., 1997; Carroll et al., 2006). Life-style of herbivores, such as aggregating behavior are aspects that are likely to be part of the attractiveness of damaged plants and of their neighbors (Bolter et al., 1997). The search for mating may explain such attraction for damaged plants by conspecifics (Kalberer et al., 2001; Arab et al., 2007). In case of uncertainty of feeding, a choice on a sub-optimal but detectable plant may also be the better choice for a slow-foraging herbivore (Carroll et al., 2006).

Second, interactions between two individuals of *Arabidopsis thaliana* can also be maladaptive or addressed to other species than snails. To our knowledge, only one case of increased attraction to herbivores in response to plant-plant signaling has been shown (Zhang et al., 2019). In this example, tomato fruits infested by the whitefly *Bemisia tabacito* expressed a defense against pathogens instead of herbivores. Neighbors of such manipulated plants also expressed unappropriated defenses and thus were more attractive to whiteflies (Zhang et al., 2019). In addition, recent studies on *Solidago sp*. showed that the volatiles emitted consecutively to herbivore damages, and their effects on neighbors was highly specialized to herbivore species (Moreira et al., 2018; Moreira and Abdala-Roberts, 2019). As we simulated herbivore attack by clipping the leaves of *A. thaliana*, induced defenses may not be specifically addressed to snails. Numerous studies showed that herbivore-induced volatiles emission attracted parasitoid insects, representing thus indirect defenses to herbivores (Turlings et al., 1990; Kessler and Baldwin, 2001; Baldwin et al., 2002; Girling et al., 2006; Dicke and Baldwin, 2010; Turlings and Erb, 2018). Girling et al. (2006) showed that attack of peach-potato aphid induced volatiles production in *A. thaliana* and attracted the aphid parasitoid *Diaeretiella rapae*. Similarly, van Poecke et al (2001) showed that *Pieris rapa* damages on *A. thaliana* led to the attraction of the parasitoid *Cotesia rubecula*. Then, *A. thaliana* may express here an olfactive signals attracting both snails and natural enemies of snails, such as dipter parasitoid (Coupland and Barker, 2004).

Studies on plant-plant interactions in response to herbivory often used field-based approaches (Fowler and Lawton, 1985; Karban et al., 2000, 2003, 2016; Karban and Baxter, 2001; Pearse et al., 2013; Morrell and Kessler, 2017; Benevenuto et al., 2020). Such an approach was associated with a dependence between the environment of the emitter and of the receiver. In other words, the signals emitted by the damaged plants may directly repel herbivores, providing indirect defenses to neighbors (as discussed in Karban et al., 2014). Here, we spatially and temporally separated plant-plant signaling of plant-herbivore interactions. The observed effects of plant-plant signaling on plant-herbivore interaction is then decorrelated of the defense produced by the emitter plants to defend itself against herbivores. More generally, if we demonstrated an effect of a prior contact on the foraging activity of snails, the underlying mechanisms, such as volatiles emission or toxin production, remain to be described.

A key aspect of modern biology is to understand how complex phenotypes are genetically controlled (Francisco et al., 2016). Genome-wide association studies are powerful and fast tools for gene hunting (Brachi et al., 2015; Francisco et al., 2016; Gloss et al., 2017). The completely-sequenced genomes of thousands of natural accessions of *A. thaliana* is then the best material for such genotype-to-phenotype studies (1001 Genomes Consortium, 2016). Here, we found that the effect of previous plant exposition to a damaged neighbor on snail behavior relied on several single-nucleotid polymorphisms (SNPs) along the genome. Most of them were associated with regions with unknown functions, thus opening the way to the exploration of new pathways involved in plant defenses (**Supplementary Tab. S1, S2, S3, and S4**). Other significant SNPs were related to plant ability to cope with stress. For instance, AT2G43110 is involved in the regulation of drought tolerance (Noman et al., 2019). AT3G03380 increases stress protection of the photosystem II in *Arabidopsis thaliana* (Luciński et al., 2011). Interestingly, we found that variation in leaf consumption was associated with the AT3G10050 gene, which encodes the first enzyme in the synthetic pathway of isoleucine, an activator of jasmonic acid (Staswick and Tiryaki, 2004) that is a well-known molecule involved in plant-plant and plant-herbivore interactions (Ray et al., 2019). Although these genes remain to be validated, for instance using mutant phenotype expressions, they are promising candidates to progress our understanding of the mechanisms involved in plant defenses against herbivore mediated by plant-plant interactions. If validated, such genetic variation underlying plant-plant interactions for induced defense against herbivore may rise possibilities for plant breeding in an agricultural context. For example, the push-pull strategy used in hundreds of sub-Saharan fields, consisting in mixing attractive and repulsive species within a field, may enhanced both ‘pull’ and ‘push’ compartments with artificial selection on alleles of interest (Bruce, 2010; Guerrieri, 2016; Pickett and Khan, 2016; Turlings and Erb, 2018). A breeding for such plant-plant interactions within species may also enhanced yield stability in varietal mixtures (Pélissier et al., 2021).

How plant-plant interactions could evolve in response to herbivory is highly debated in ecology (Dicke and Baldwin, 2010; Heil and Adame-Álvarez, 2010; Heil and Karban, 2010; Heil, 2014; Karban et al., 2014; Karban, 2021). Discussions were substantial on the advantage for an attacked plant to produce a signal that prevent and help potential competitor (Agrawal, 2000; Heil and Adame-Álvarez, 2010; Heil and Karban, 2010). At the opposite, the advantages for a plant receptor of such signal was often considered as adaptive, in the case of increased resistance to herbivory after being in contact with a damaged plant. Theoretical expectations on the evolution of an increased attractivity to herbivores when receiving cues from attacked plants are missing in the literature (as discussed in Pearse and Karban, 2013). Encouraging continuous movement of herbivores between attractive plants may limit biomass consumption at the group level (Morrell and Kessler, 2017). We propose that mutual benefits across close relatives could arise if increasing attractivity of plant neighbors dilutes the effect of the herbivore at the neighborhood level.

## Acknowledgements

We are very grateful to Ana Elkhaïm and Maëva Tremblay for their help on measurements. We also thank the technical platform “Plateforme des Terrains d’Expérience du LabEx CeMEB,” in Montpellier.

## Author Contribution

AE, FV, CV, and DV designed the experiment. AE, FV, DV, CV, AH, and TM performed the experiment. AE, AH, TM, and PM measured response variables from films and pictures. AE analyzed the data and wrote the article with comments from all authors.

## Data availability

The data that support the findings of this study are openly available in Data.INRAE at https://doi.org/10.57745/QN7MMM.

## Supplementary

**Figure S1**. Leaf index consumption.

**Figure S2**. Distribution of first duration spent on plants across natural accessions of *Arabidopsis thaliana*.

**Video S1**. Example of foraging activity of a snail in an arena on two plants of the “6830” genotype.

**Table S1, 2, 3, and 4**. List of single nucleotid polymorphisms associated with the first choice, first duration, proportion of time and leaf consumption index, respectively.

**Table S5**. Effects of minor alleles of significant SNPs on response variables.

**Supplementary 1**. Attractivity of clipped plants.

